# High-density genome-wide association study points out major candidate genes for resistance to infectious pancreatic necrosis in rainbow trout

**DOI:** 10.1101/2025.04.19.649654

**Authors:** Jonathan D’Ambrosio, Yoannah François, Thierry Morin, Sébastien Courant, Alexandre Desgranges, Pierrick Haffray, Bertrand Collet, Pierre Boudinot, Florence Phocas

## Abstract

**Background:** This study focuses on genetic resistance to infectious pancreatic necrosis (IPN), a highly contagious disease caused by an aquatic birnavirus (IPNV) which especially affects salmonids worldwide. The objectives were to estimate the heritability of IPN resistance and to fine map quantitative trait loci (QTL) using a Bayesian Sparse Linear Mixed Model to identify candidate genes possibly linked to IPN resistance in two successive generations from a French commercial strain of rainbow trout. For each generation, 2,000 fish were experimentally exposed by bath to IPNV and mortalities were monitored daily during 5 weeks. All fish were genotyped using a medium-density 57K SNP chip and imputed to high-density genotypes (665K SNP).

**Results:** The mean survival rate was 70% after 37 days, with a higher survival rate in the second generation compared to the first one (78% versus 61%). Heritability was moderate (∼0.20). Approximately 74% of the genetic variance of IPN resistance was explained by a few tens of SNP. In total, 25 QTL were mapped on 10 chromosomes. Among them, 7 were detected with very strong evidence on chromosomes 1, 14, 16 and 28. The most interesting QTL were associated to top SNP with mean survival rate differences over 20% between the beneficial and detrimental homozygous genotypes. Those SNP were all located within promising functional candidate genes on chromosome 1 (*uts2d*, *rc3h1*, *ga45b*) and chromosome 16 (*irf2bp*, *eif2ak2)*, all these genes being associated to the regulation of inflammatory pathways. A key factor of the genetic differences in susceptibility to IPNV among fish is PKR, the dsRNA-dependent serine/threonine-protein kinase encoded by the *eif2ak2* gene.

**Conclusions:** All genes associated to the most significant QTL on chromosomes 1 and 16 are involved in the regulation of inflammatory pathways, strongly suggesting a central role of inflammation in the IPN resistance in rainbow trout. These findings offer the possibility of marker-assisted selection for rapid dissemination of genetic improvement for IPN resistance.

## Background

Infectious pancreatic necrosis (IPN) is a severe viral disease that affects salmonids worldwide. In rainbow trout farming, IPN was first described in USA in 1940 [1] and reported in 1964 with catastrophic fry losses in French farms [2]. Its etiological agent, IPNV, is a double-stranded bi-segmented non-enveloped RNA birnavirus belonging to the Aquabirnavirus genus [3]. The two genomic segments, called A and B, encodes five viral polypeptides (VPs): VP1, an RNA-dependent RNA polymerase enzyme (segment B); VP2, the major capsid protein; VP3, an internal/minor capsid protein; VP4, a serine-lysine protease; and VP5, a non-structural protein (segment A). According to a phylogenetic classification based on VP2, seven genogroups are defined to date [4]. IPNV, which infects over 63 different species of fish, molluscs, and crustaceans, is highly prevalent in both farmed and wild fish [5]. In Europe, where a large number of countries cultivate salmonids, the majority of the farm isolates belongs to genogroup 5 [4]. Mainly fry and juveniles less than six months old are affected by the disease with up to 90% mortality on fry at start-feeding. The main symptoms are unusual behavior (abnormal swimming, i.e spinning), skin melanism, abdominal distension, catarrhal lesions, and necrosis of the exocrine pancreas and liver tissues [5]. However, this highly contagious disease also affects farmed salmon at the post-smolt stage [3]. Survivors of an infection may become healthy adult carriers that can infect naïve fish [3]. The disease spreads horizontally via infected water and fish, but can also be transmitted vertically through eggs. Currently, no effective treatment is available. Several vaccine candidates (DNA, subunit, attenuated, inactivated, recombinant) were developed and tested, and some commercial vaccines are available but it remains difficult to induce effective immune protection in the early stages most affected by the virus [6–8]. However, good husbandry practices, such as maintaining high water quality and low stocking density, and avoiding mixing of batches, help to reduce disease incidence [3].

Selective breeding is an interesting alternative to control infectious diseases in salmonids [9]. Several studies reported a significant genetic variation for resistance to IPN in Atlantic salmon with moderate narrow-sense heritability estimates ranging from 0.31 to 0.45 [10]. In rainbow trout, the estimates of heritability are few for IPN resistance with values varying from 0.24 to 0.39 on the observed survival scale, depending on either genomic or pedigree information was used to derive the estimates in a Chilean population [10,11], and at 0.30 in a Norwegian breeding strain from AquaGen [12]. In Atlantic salmon, two research groups [13,14] independently discovered a major QTL for IPN resistance on chromosome 26, using a few large full-sib fish groups coming from a Scottish and a Norwegian breeding population, respectively. The QTL turned out to be responsible for more than 80% of the genetic variation in IPN resistance both at the fry and the post-smolt life stages in Atlantic salmon [15,16]. This QTL for IPN resistance was an ideal case for marker-assisted selection (MAS) programs in Atlantic salmon, as it explained a large fraction of the phenotypic variance of a trait with high economic value, and was still segregating with an intermediate allele frequency in many populations, the high-resistance allele being partly dominant over the low-resistance allele [17]. In Norway, MAS for IPN resistance has contributed to a 75 % decline in the number of IPN-outbreaks within a few years from 2009 to 2013 [18]. A functional mutation, underlying this QTL, was identified in the *epithelial cadherin* gene (*cdh1*). In a co-immunoprecipitation assay, *cdh1* found to bind to IPNV virions, strongly indicating that the protein was involved in the internalization of the virus [16]. However, a new variant of IPNV has been recently detected in Norway with mutations that have made the virus capable to cause a disease even in the genetically IPN resistant fish for the major QTL [19].

In rainbow trout, the few QTL identified in the literature were not located in the vicinity of the salmon major QTL. The use of low-resolution molecular markers has shown the presence of two genomic regions associated with resistance and susceptibility to IPN [20,21]. In two families, one hybrid family between a IPN resistant and a IPN susceptible Japanese rainbow strains and a backcross family, two QTL were found on linkage groups LG 3 and LG 22 [20,21], corresponding respectively to chromosomes 14 and 16 [22]. These two QTL are on linkage groups that have no clear homology with the linkage groups in which QTL were identified for IPN resistance in Atlantic salmon [23,24]. The first medium-density GWAS using SNP markers on a small set of 721 phenotyped and genotyped Chilean rainbow trout from 58 full-sib families [11] provided some moderate evidence for QTL mainly on chromosomes 5, 13, 21 and 23. Although the *cdh1* gene is located on chromosome 5 in rainbow trout, the QTL identified on this long chromosome was located at the opposite end. In addition, based on a batch-challenge test and 57K SNP genotyping of 1723 rainbow trout from 46 full-sib families, a US patent was deposited by AquaGen AS [25] revealing a significant IPN QTL region in chromosome 1 with the 3 most significant SNPs being located between 16 and 19 Mb on chromosome 1 of the Arlee reference genome [26].

With a high-density SNP chip now being available in rainbow trout [27], the objectives of this research were to estimate the heritability of IPN resistance and to perform QTL fine mapping to identify candidate genes possibly linked to IPN resistance in two successive generations from a French commercial strain of rainbow trout.

## Methods

### Animals

The fish challenged were produced by parents from the 8^th^ (G8) and the 9^th^ (G9) generations of selection of a commercial breeding program developed by Milin Nevez breeding company (Plouigneau, France). For G8, the experimental stock was established from 81 dams and 91 sex-reversed neomales with 10 independent full-factorial mating designs (∼8 dams × 8-10 neomales in each factorial). For G9, the experimental stock was established from 90 dams and 98 neomales with 10 independent full-factorial mating designs (9 dams × 8-10 neomales in each factorial).

A batch of eyed eggs was transferred to the SYSAAF-ANSES FORTIOR Genetics platform (Plouzané, France) at about 200-250 degree-days. Fry were reared in tanks in an opened flow-through system with filtered freshwater at 10°C ±2 temperature. About 15 to 20 days after start feeding (about 700 degree-days, or 1,5g), fry were transferred to adapted tanks for the infectious challenges. Fish were individually identified post-challenge using the DNA barcode of the individual fin sampling.

### Challenge tests

Fish were challenged in year 2018 for G8 and in 2020 for G9, respectively. Both infectious challenges were conducted the same way. Each generation, 2000 fry were exposed to a strain of IPNV isolated from affected farm during a bath in static aerated freshwater at 10°C ±2 containing 1.10^5^ Tissue Culture Infectious Dose (TCID) _50_/ml of virus. For G8 and G9, the strains used were identified NN193 (isolated in 2015) and NPI11125 (isolated in 2019), respectively, both belonging to genogroup 5. After 3 hours, water was restarted in open circuit, with a minimum hourly renewal, at 10°C ±2. Challenges lasted respectively 37 and 36 days post-infection (dpi), during which fish were monitored daily and fed at least twice a day. Days of mortality were recorded and DNA samples were individually collected for all dead fish (caudal fin sampled and stored in alcohol tubes in 4°C). At the end of the challenges, survivors were sacrificed using a lethal dose of Eugenol (180 ppm; Fili@Vet Réseau Cristal) and their DNA samples were also collected. To control the sanitary status of infected and uninfected (control) fish, broad virological (NRL for regulated fish diseases) and bacteriological (Labocea, Quimper) analyses were performed at the mortality peak on both groups. IPNV detection was carried out according to the AFNOR UN 47-222 standard by culture on EPC (epithelioma papulosum cyprinid), BF2 (Bluegill fry) or CHSE (Chinook salmon embryo) cell lines then seroneutralization.

### Genotyping and Imputation

Fin samples from 1878 (G8) and 1992 (G9) challenged fish and their 372 parents were sent to the INRAE genotyping platform Gentyane (Clermont-Ferrand, France) for DNA extraction and genotyping. The 3870 challenged fish as well as the 186 parents of G9 were genotyped for 57,501 SNPs using the medium-density (MD) Rainbow Trout Axiom® 57K SNP array from Thermo Fisher (Palti et al., 2015).

The 186 parents of G8 as well as 95 sibs of G9 challenged fish were genotyped for 664,531 SNPs using the high-density (HD) Rainbow Trout Axiom® 665K SNP array [27]. Then, SNPs with probe polymorphism and multiple locations on the Arlee reference genome assembly (GCA_013265735.3; [26]) were discarded as described in [27].

For both MD and HD genotypes, PLINK v1.9 software [28,29] was used for keeping only SNPs with deviation from Hardy-Weinberg equilibrium with a p-value > 0.000001, a minor allele frequency greater or equal to 5%, and both SNP and sample call rates above 98%. After quality control (QC), 409,786 SNPs and 27,130 SNP were retained for HD and MD genotypes, respectively. In total 162 and 2 fish samples did not pass QC respectively for MD and HD genotypes.

Parentage assignment was done using 1,000 randomly sampled markers with the R package APIS [30,31] with a positive assignment error rate set to 1%.

The imputation of the MD genotypes into HD genotypes for the 3707 offspring (1757 G8, 1950 G9) was run using FIMPUTE3 software [32] utilizing quality-filtered genotypes and pedigree information from parents. The correctness of imputation was checked by mendelian error testing. In average there were 6 mendelian errors per SNP with a maximum of 3 errors (< 1% of the progeny) observed for 75% of the 409,786 SNPs. A last quality filter was used to remove SNPs with over 100 mendelian errors after imputation. In total 394,101 SNPs were finally retained for the analysis of HD genotypes, with 3 mendelian observed in average per SNP and a maximum of 2 errors for 75% of the SNPs.

The final dataset for the genomic analysis contained 3707 phenotyped progeny with imputed genotypes for 394,101 SNPs.

### Estimation of genetic parameters

Variance components for IPN resistance were estimated using the restricted maximum likelihood method applied to a (G)BLUP linear animal model and AIREML algorithm in BLUPF90 software [33]. The cohort effect was the only fixed effect included in the model. In total, 44,367 animals were related through the pedigree relationship matrix for PBLUP evaluation, tracing back 9 generations of ancestors of the 3855 phenotyped animals. For GBLUP evaluation, the pedigree relationship matrix was replaced by a genomic relationship matrix [34].

### QTL mapping and credibility intervals

The genome-wide association study (GWAS) was based on a Bayesian Sparse Linear Mixed Model (BSLMM) that assumes that all SNPs have at least a relatively small effect but also that some SNPs may have a large effect [35]. The BSLMM was applied on the IPN challenge resistance phenotypes (here, a binary trait corresponding to the status ‘dead’ or ‘alive’), corrected by the cohort effects estimated under a linear model. The resulting residues were integrated as the vector of phenotypes **y** described as follows:

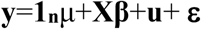

where **1_n_** is an n-vector of 1s, µ is a scalar representing the phenotype mean, **X** is an n × p matrix of genotypes measured on n individuals at p SNPs, **β** is the corresponding p-vector of the SNP effects; **u** is a vector of random additive genetic effects distributed according to *N*(0, **K**σ²_b_), with σ²_b_ the additive genetic variance and **K** the genomic relationship matrix; and **ε** is a n-vector of residuals *N*(0, **I** σ²_e_), σ²_e_ is the variance of the residual errors.

Assuming **K** = **XX^T^** /p, the SNP effect sizes can be decomposed into two parts: α that captures the small effects that all SNPs have, and β that captures the additional effects of some large effect SNPs. In this case, u = Xα can be viewed as the combined effect of all small effects, and the total effect size for a given SNP is **γ**_i_ = α_i_ + β_i_. The individual SNP effects **γ**_i_ are sampled from a mixture of two normal distributions, **γ**_I_ ∼ π N(0, σ²_a_+σ²_b_) + (1−π) N(0,σ²_b_) where σ²_b_ is the variance of small additive genetic effects, σ²_a_ is the additional variance associated to large effects and π is the proportion of SNPs with large effects.

The BSLMM was implemented in our work using the Genome-Wide Efficient Mixed Model Association (GEMMA) software based on a Markov chain Monte Carlo (MCMC) method applied to a linear mixed model (‘-bslmm 1’ option) on our binary trait as proposed by Zhou et al [35] for survival trait. A total of 2.2 million iterations (-s option) were performed with a burn-in of 200,000cycles and record one state in every 100 iterations for further analysis. In addition, to ensure convergence of the distribution of the hyper-parameter π, the minimum and maximum numbers of SNPs that was sampled to be included in the model were set to 1 (-smin option) and 300 (-smax option), respectively. We ran 3 chains MCMC with 3 different initial random seeds in order to check the convergence of the estimates.

The MCMC sampling of all parameters values from the posterior distribution allows to estimate all the SNP effects 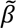, but also the hyper-parameter π and the posterior inclusion probability (PIP) of each SNP that quantifies the proportion of samples in which the SNP has a large effect in the model. This PIP value indicates the strength of the evidence that the SNP has to be included in the model, and can therefore be used for QTL mapping.

To define a minimum threshold for the strength of the evidence for a given SNP, Stephens and Balding [36] have proposed to calculate the Bayesian Factor BF = [PIP/(1-PIP)] / [ π/(1-π)]. The logBF was computed as twice the natural logarithm of the BF to be in the usual range of likelihood ratio test values, and a minimum threshold value of logBF=10 (BF ≈ 150) was used for defining strong evidence for a QTL [37]. A more stringent value of logBF=12 (i.e. BF ≈ 400) was also considered for pointing out very strong evidence for a QTL [38].

The GWAS results were visualized via a Manhattan plot where all negative values for logBF were set to 0. The GWAS results were visualized via a Manhattan plot where all negative values for logBF were set to 0.

To account for differences in allele frequencies and linkage disequilibrium between SNPs, credibility intervals were determined by including any SNP in a QTL region as soon as a SNP had a logBF ≥ 7 within a 100 kb sliding window from the top SNP with evidence for the QTL (i.e. logBF ≥ 10).

### Effects of QTL genotypes

To assess how the QTL genotypes affect the survival rate, the effects of top SNPs in each QTL region were analyzed. To do so, we adjusted the survival rates observed by their cohort effect in order to be able to compare the two challenge test results. Then, we averaged these survival rates across the two cohorts to facilitate the discussion of the results.

### Gene annotation

Genes within QTL regions were annotated using the NCBI O. mykiss Arlee genome assembly USDA_OmykA_1.1. (GCA_013265735.3,[26]). In addition to NCBI gene summaries, functional information for genes was extracted from the human gene database GeneCards® (https://www.genecards.org/) that also includes protein summaries from UniProtKB/Swiss-Prot (https://www.uniprot.org/uniprotkb/).

## Results

### Challenge tests

The mean survival rate was 70% after 37 days, with a higher survival rate in G9 compared to G8 (78% versus 61%). Mortality kinetics were relatively similar between the two cohorts, but with a more rapid decline in survival between 8 and 15 days for G8 (Fig. 1).

**Figure 1.**
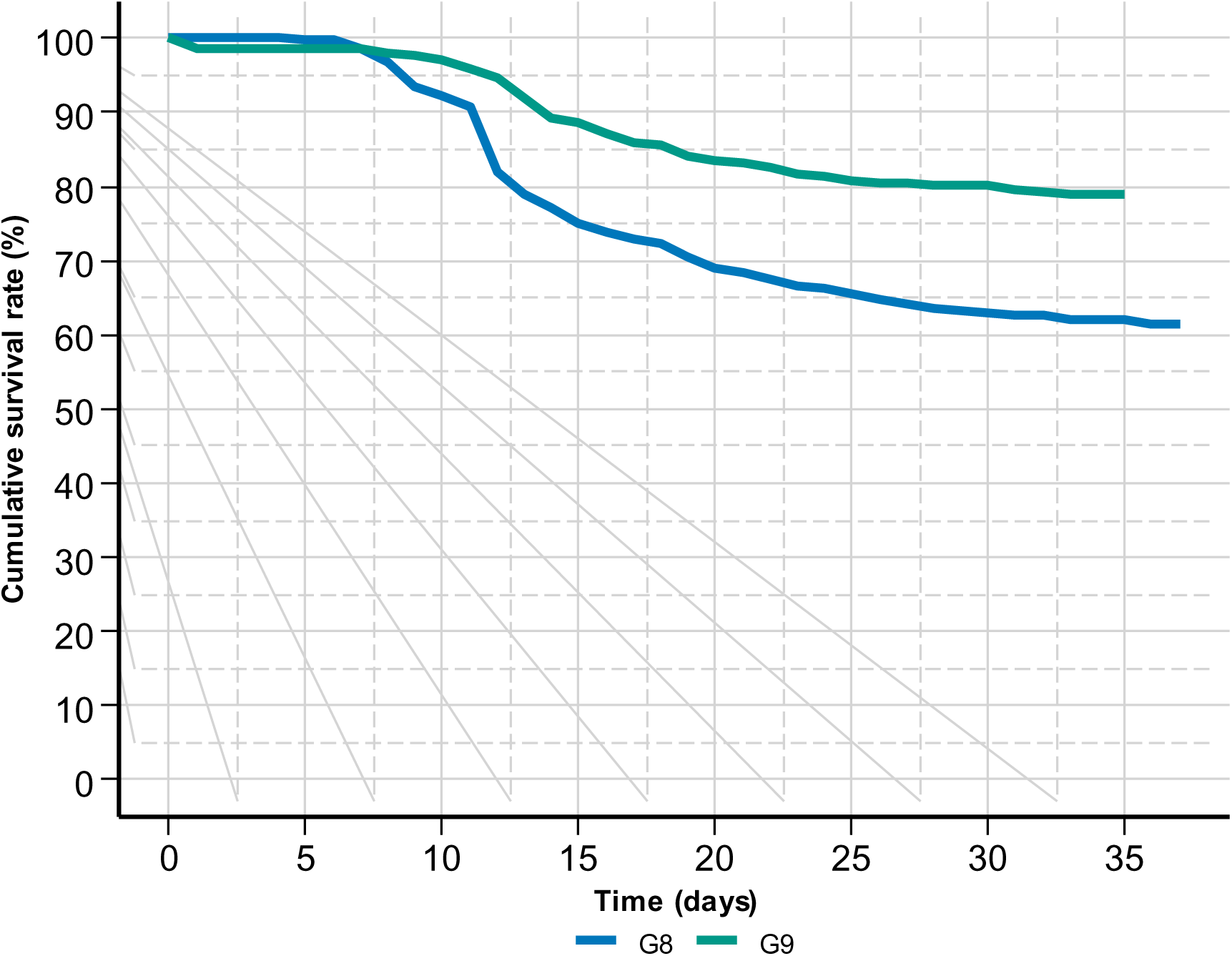
Evolution of the Kaplan-Meier cumulative survival rate during the challenge tests of the two cohorts.

**Figure 2.**
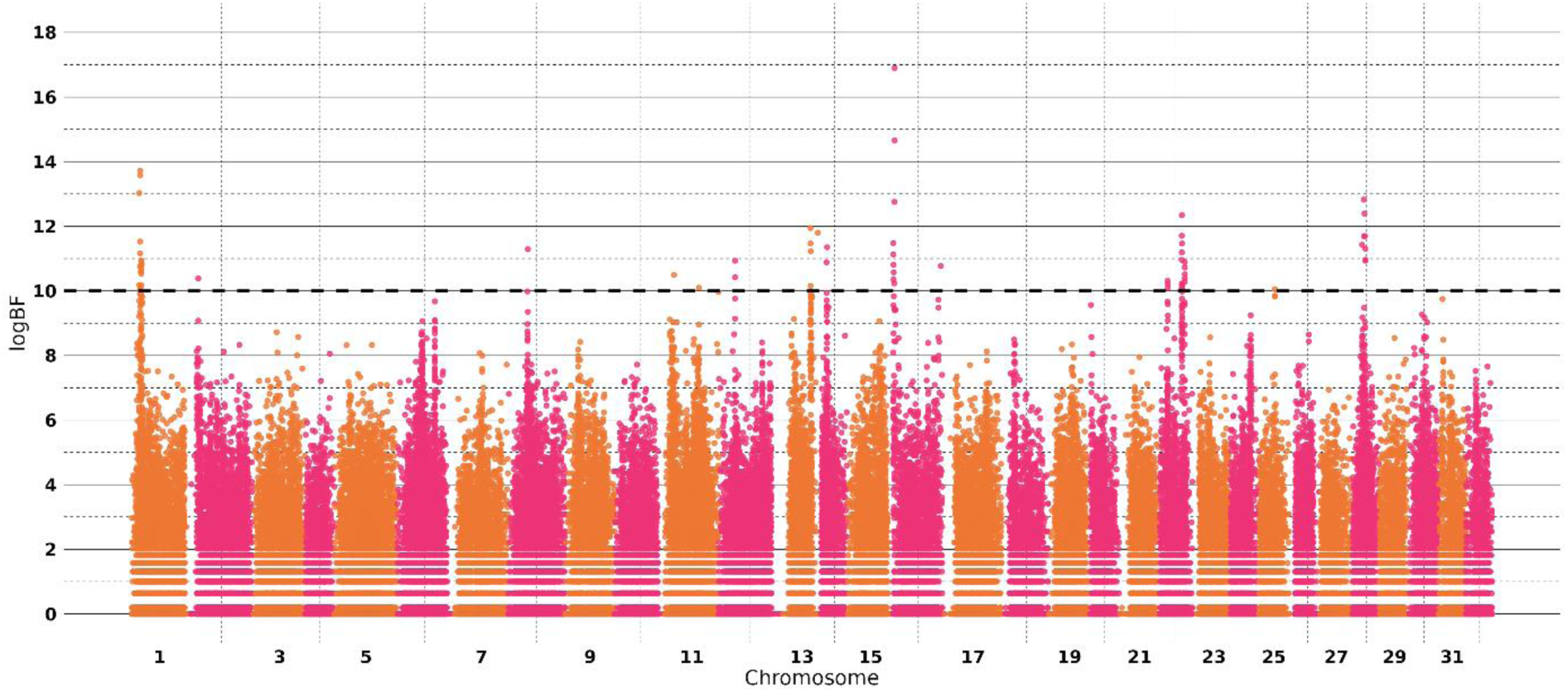
Manhattan plot showing associations between SNPs and IPN resistance. The dashed line represents a threshold value of logBF=10, indicating a strong association between SNP and IPN resistance.

The presence of IPNV was confirmed in dead fry following experimental infections by observation of cytopathic effects in cell cultures and identification by seroneutralization. Bacteriological analyses did not reveal any pathogenic germs. No infectious agent was detected in uninfected control animals.

### Genetic architecture of IPN resistance

Heritability estimates of survival were very moderate, close to 0.20 on the observed scale for both BLUP and GBLUP models (Table 1). The corresponding parameter (PVE) under BSLMM was estimated at a slightly lower value that was consistent across the 3 MCMC runs. Across the 3 MCMC runs of BSLMM, approximately 74% of the total genetic variance of IPN resistance was explained by a moderate number of SNPs with a median value of 63 SNPs fitted with a large effect in the model.

**Table 1.** Heritability estimates for IPN resistance on the observed scale according to the genetic model.

| Model – heritability symbol | PBLUP – $h^2$ | GBLUP – $h^2$ | BSLMM - PVE |
| --- | --- | --- | --- |
| Mean estimate (±SE) | 0.206 (±0.005) | 0.190 (±0.023) | 0.154 (±0.018) |

Major evidence (logBF ranging from 14.7 to 16.9 for two SNPs) was found on chromosome 16 for a QTL (Q16.3) harbouring the region of eukaryotic translation initiation factor 2-alpha *kinase 2* (*eif2ak2*) gene. This gene, also known as *pkr (protein kinase RNA-activated*), is a conserved interferon stimulated gene that mediates virus induced protein shut off via inhibition of translation, promotes apoptosis, fosters type I IFN responses induced by some viruses and modulates inflammation through activation of MAPK and NFkB pathways [39]. At the two top SNPs located within *pkr* (Table 2), the same major allele G correspond to the resistant allele (R) and the minor allele A is the susceptible allele (S) with a MAF of 15.9%. Of note, the major allele G for the first top SNP Affx-1237475214 is the Arlee alternative allele while the major allele G for the second top SNP Affx-1248466717 is the Arlee reference allele (see Additional file 1: Table S1).

**Table 2.**
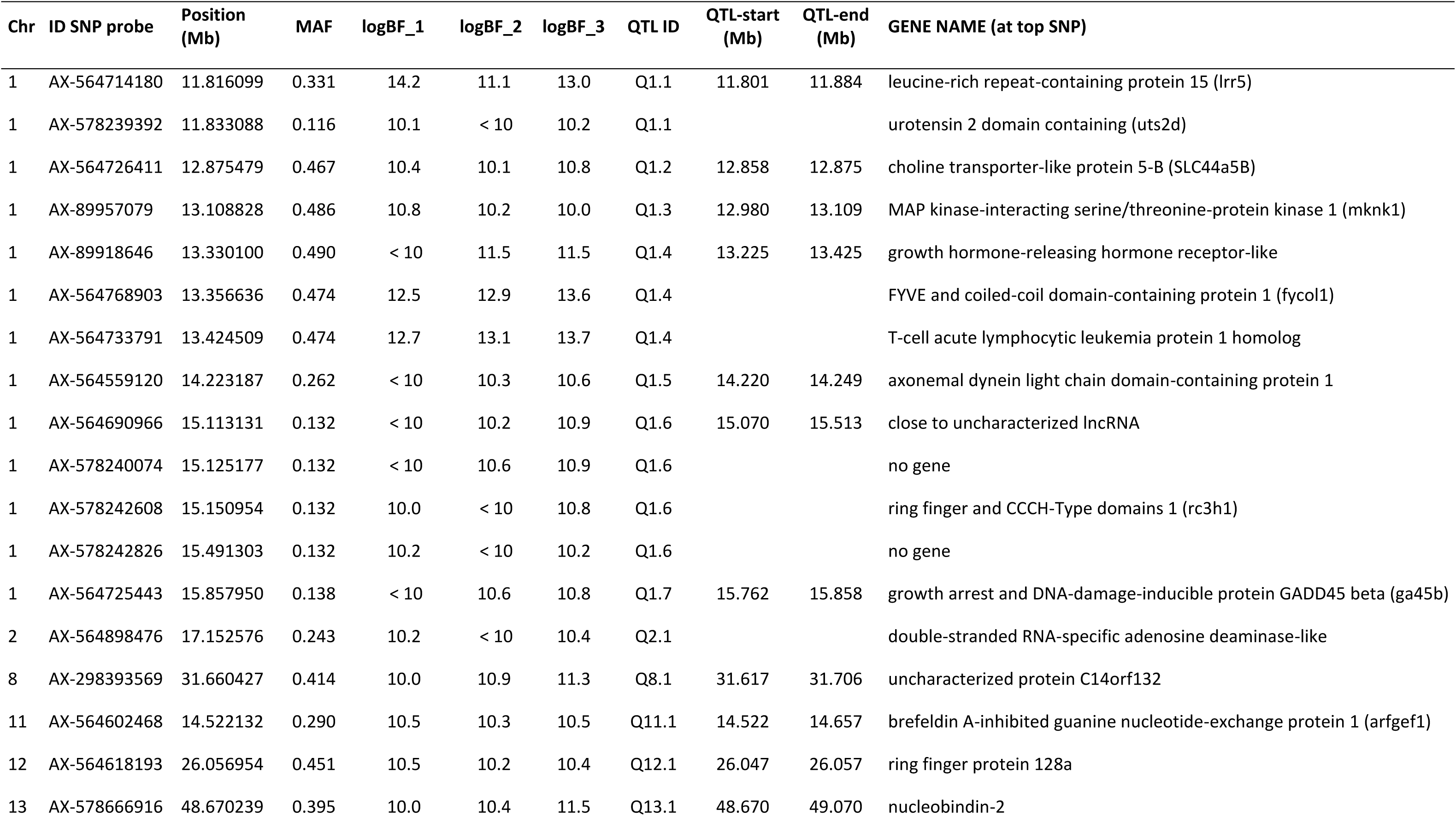
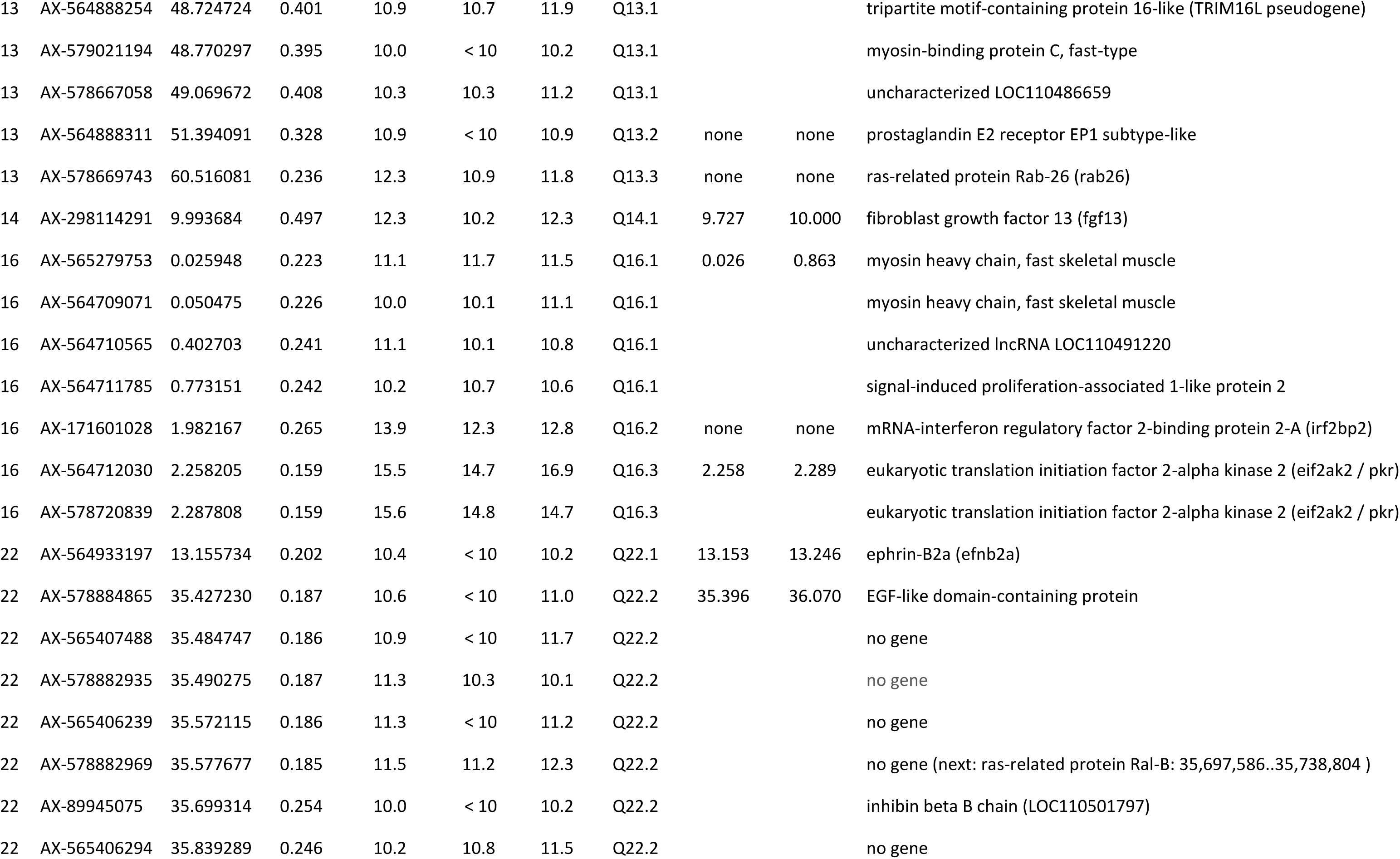
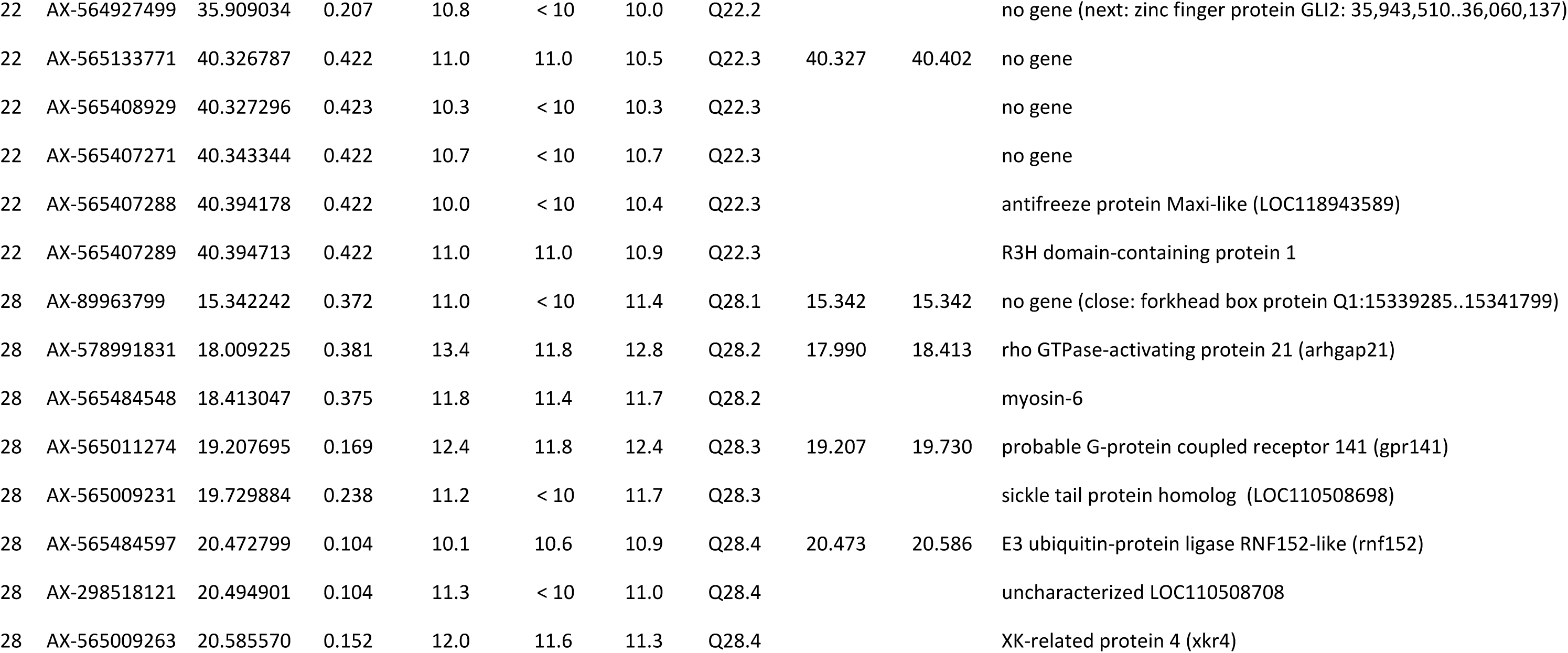
Top SNPs, Bayes Factor statistics, credibility intervals and candidate genes for IPN resistance in French rainbow trout.

Six other QTL were detected on chromosomes 1 (Q1.1, Q1.4), 14 (Q14.1), 16 (Q16.2), and 28 (Q28.2, Q28.3) with very strong evidence (logBF > 12 for at least 2 out of the 3 MCMC runs). Some top SNPs for all these six QTL were also identified within genes (Table 2).

Several genes associated to QTL are orthologous to genes controlling either the type I IFN response and/or inflammation (*uts2d* and *rc3h1* on chr 1; *rab26* on chr 13; *irf2bp2* located in Q16.2 close to *pkr* on chr 16; *gpr141* and *rnf52*, respectively in Q28.3 and Q28.4 on chr 28). Located in Q1.1, *uts2d* is a homolog of the mammalian *urotensin-2* that induces a number of pro-inflammatory genes including TNF-α, IL-1β, IFN-γ, IL-8 and leukotriene C4, or activates pathways like NFκB and IRF3 signaling [40]. In Q1.6, the gene *rc3h1* (also named *roquin-1*) encodes for an anti-inflammatory and regulatory factor [41,42].

The other genes associated to QTL played various roles in the biology of the cell, which might be connected to the virus cycle or to antiviral processes: choline/phospholipid metabolism (s*lc44a5b* in Q1.2; *xkr4* in Q28.4), translation control (*mknk1* in Q1.3), intracellular vesicular trafficking (*fyco1* in Q1.4; *argef1* in Q11.1), and actine remodelling (*arhgap21* in Q28.2 on chr 28).

### QTL effects on IPN resistance

Survival rates were calculated from all the 53 top SNPs in each of the two cohorts. The average differences in survival rate between the two homozygous genotypes as well as between the major homozygote and the heterozygote were derived and presented in Table 3.

**Table 3.** Survival rates differences according to the genotypes and their frequency at the top SNPs for resistance to IPNV.

| Chr | ID SNP probe | position (Mb) | Major allele | Minor allele | QTL ID | Difference in % survival between major homozygotes and minor homozygotes | Difference in % survival between major homozygotes and heterozygotes | % major homozygotes | % minor homozygotes |
| --- | --- | --- | --- | --- | --- | --- | --- | --- | --- |
| 1 | AX-564714180 | 11,816099 | C | T | Q1.1 | 13.9 | 3.7 | 0.34 | 0.18 |
| 1 | AX-578239392 | 11,833088 | G | A | Q1.1 | 25.5 | 8.7 | 0.74 | 0.02 |
| 1 | AX-564726411 | 12,875479 | C | A | Q1.2 | -12.0 | -14.3 | 0.42 | 0.13 |
| 1 | AX-89957079 | 13,108828 | A | G | Q1.3 | 13.7 | 0.1 | 0.16 | 0.35 |
| 1 | AX-89918646 | 13,330100 | T | G | Q1.4 | 13.7 | 0.1 | 0.16 | 0.36 |
| 1 | AX-564768903 | 13,356636 | A | C | Q1.4 | 15.5 | 2.6 | 0.18 | 0.33 |
| 1 | AX-564733791 | 13,424509 | G | T | Q1.4 | 15.7 | 2.5 | 0.18 | 0.33 |
| 1 | AX-564559120 | 14,223187 | C | T | Q1.5 | 17.3 | 5.0 | 0.50 | 0.09 |
| 1 | AX-564690966 | 15,113131 | G | A | Q1.6 | 29.8 | 6.8 | 0.72 | 0.02 |
| 1 | AX-578240074 | 15,125177 | C | A | Q1.6 | 29.8 | 6.8 | 0.72 | 0.02 |
| 1 | AX-578242608 | 15,150954 | G | T | Q1.6 | 29.8 | 6.9 | 0.72 | 0.02 |
| 1 | AX-578242826 | 15,491303 | C | A | Q1.6 | 27.4 | 7.4 | 0.72 | 0.02 |
| 1 | AX-564725443 | 15,857950 | G | A | Q1.7 | 30.0 | 6.5 | 0.71 | 0.02 |
| 2 | AX-564898476 | 17,152576 | C | T | Q2.1 | 11.4 | 4.4 | 0.55 | 0.06 |
| 8 | AX-298393569 | 31,660427 | A | C | Q8.1 | -8.9 | -5.9 | 0.37 | 0.15 |
| 11 | AX-564602468 | 14,522132 | G | A | Q11.1 | -13.0 | -6.0 | 0.56 | 0.06 |
| 12 | AX-564618193 | 26,056954 | C | T | Q12.1 | 7.3 | 5.2 | 0.30 | 0.19 |
| 13 | AX-578666916 | 48,670239 | A | G | Q13.1 | 8.5 | 6.7 | 0.32 | 0.17 |
| 13 | AX-564888254 | 48,724724 | T | C | Q13.1 | 8.3 | 6.8 | 0.31 | 0.17 |
| 13 | AX-579021194 | 48,770297 | C | A | Q13.1 | 8.3 | 6.8 | 0.32 | 0.17 |
| 13 | AX-578667058 | 49,069672 | G | A | Q13.1 | 8.6 | 6.0 | 0.30 | 0.19 |
| 13 | AX-564888311 | 51,394091 | G | T | Q13.2 | 8.5 | 7.6 | 0.38 | 0.13 |
| 13 | AX-578669743 | 60,516081 | T | G | Q13.3 | 6.8 | 4.4 | 0.50 | 0.07 |
| 14 | AX-298114291 | 9,993684 | C | A | Q14.1 | 12.3 | 6.2 | 0.24 | 0.26 |
| 16 | AX-565279753 | 0,025948 | A | C | Q16.1 | 19.2 | 10.5 | 0.54 | 0.07 |
| 16 | AX-564709071 | 0,050475 | G | T | Q16.1 | 19.0 | 10.4 | 0.54 | 0.07 |
| 16 | AX-564710565 | 0,402703 | T | C | Q16.1 | 18.5 | 9.3 | 0.49 | 0.08 |
| 16 | AX-564711785 | 0,773151 | G | A | Q16.1 | 18.5 | 9.4 | 0.49 | 0.08 |
| 16 | AX-171601028 | 1,982167 | T | C | Q16.2 | 19.6 | 10.6 | 0.49 | 0.10 |
| 16 | AX-564712030 | 2,258205 | G | A | Q16.3 | 24.1 | 12.4 | 0.64 | 0.04 |
| 16 | AX-578720839 | 2,287808 | G | A | Q16.3 | 24.1 | 12.4 | 0.64 | 0.04 |
| 22 | AX-564933197 | 13,155734 | C | T | Q22.1 | -9.7 | -6.4 | 0.63 | 0.04 |
| 22 | AX-578884865 | 35,427230 | C | T | Q22.2 | 12.3 | 7.9 | 0.64 | 0.03 |
| 22 | AX-565407488 | 35,484747 | A | G | Q22.2 | 12.0 | 8.1 | 0.64 | 0.03 |
| 22 | AX-578882935 | 35,490275 | T | C | Q22.2 | 11.9 | 8.0 | 0.64 | 0.03 |
| 22 | AX-565406239 | 35,572115 | A | G | Q22.2 | 11.5 | 8.1 | 0.64 | 0.03 |
| 22 | AX-578882969 | 35,577677 | G | A | Q22.2 | 12.1 | 8.1 | 0.64 | 0.03 |
| 22 | AX-89945075 | 35,699314 | G | A | Q22.2 | 11.2 | 6.2 | 0.55 | 0.07 |
| 22 | AX-565406294 | 35,839289 | A | C | Q22.2 | 12.5 | 6.0 | 0.55 | 0.06 |
| 22 | AX-564927499 | 35,909034 | T | C | Q22.2 | 13.8 | 5.8 | 0.60 | 0.05 |
| 22 | AX-565133771 | 40,326787 | A | C | Q22.3 | 10.6 | 3.0 | 0.31 | 0.19 |
| 22 | AX-565408929 | 40,327296 | C | A | Q22.3 | 10.4 | 3.0 | 0.31 | 0.19 |
| 22 | AX-565407271 | 40,343344 | A | G | Q22.3 | 10.4 | 2.5 | 0.31 | 0.20 |
| 22 | AX-565407288 | 40,394178 | A | G | Q22.3 | 10.5 | 3.0 | 0.31 | 0.19 |
| 22 | AX-565407289 | 40,394713 | A | G | Q22.3 | 10.6 | 2.9 | 0.31 | 0.20 |
| 28 | AX-89963799 | 15,342242 | T | G | Q28.1 | -10.2 | -4.7 | 0.38 | 0.14 |
| 28 | AX-578991831 | 18,009225 | G | T | Q28.2 | 13.2 | 4.5 | 0.40 | 0.15 |
| 28 | AX-565484548 | 18,413047 | T | C | Q28.2 | 13.2 | 3.9 | 0.41 | 0.14 |
| 28 | AX-565011274 | 19,207695 | T | C | Q28.3 | -11.7 | -6.7 | 0.69 | 0.03 |
| 28 | AX-565009231 | 19,729884 | A | G | Q28.3 | -9.3 | -4.1 | 0.59 | 0.05 |
| 28 | AX-565484597 | 20,472799 | A | C | Q28.4 | -8.2 | -8.6 | 0.81 | 0.01 |
| 28 | AX-298518121 | 20,494901 | T | G | Q28.4 | -8.2 | -8.6 | 0.81 | 0.01 |
| 28 | AX-565009263 | 20,585570 | A | G | Q28.4 | -12.6 | -5.9 | 0.70 | 0.02 |

The SNPs identified as having the strongest phenotypic effect (> 24% between the two homozygotes, Table 3) were located on 3 QTL on chr 1 (Q1.1, Q1.6, Q1.7) and Q16.3 on chr 16. All the 13 SNPs identified in the 7 QTL detected on chromosome 1 exhibited at least a significant difference of 12% of survival rates between the two homozygous genotypes. These 13 SNPs also showed complete or incomplete beneficial dominance effect of the major alleles except for Q1.2 for which a complete beneficial dominance effect was observed for the minor allele of AX-564726411 (Table 3).

The two main differences (29.8% and 30.0%) in the survival rates of homozygous genotypes were observed for the SNPs AX-578242608 and AX-564725443, respectively located in the *rc3h1* gene (Q1.6) and close to *gadd45b* (*growth arrest and DNA-damage-inducible protein GADD45 beta)* gene (Q1.7). For these two SNPs, survival rates for individuals with favorable homozygous genotypes were 64% and 81% for, respectively, G8 and G9, while the rates dropped to 29% and 56% for the unfavorable homozygous genotypes (see Additional file 1: Table S2). The top SNPs across these two QTL are in very strong linkage disequilibrium (DL=0.946) (see Additional file 1: Table S3).

Regarding the 3 QTL on chromosome 16, all the 7 top SNPs exhibited at least a difference of 18.5% of survival rates between the two homozygous genotypes (Table 3) and showed pure additive genetic effects as the survival rate of the heterozygotes was very close to the mean survival rate of the two homozygotes for each of these 7 SNPs. Fish with resistant homozygous genotypes for the main QTL (Q16.3) on chromosome 16 had a survival rate of 67% in G8 and 83% in G9, compared to 38% in G8 and 63% in G9 for individuals with the susceptible homozygous genotypes (see Additional file 1: Table S2).

For all SNPs in the 15 remaining QTL regions (on chromosomes 2, 8, 11, 13, 14, 22 and 28), the differences in survival rates between homozygote genotypes were moderate, with absolute values ranging from 7 to 13%. The favorable genotypes were the major homozygous ones except for the SNPs corresponding to Q8.1, Q11.1, Q22.1, Q28.1, Q28.3 and Q28.4 for which the beneficial alleles were the minor ones (see Table 3).

The very best 2-QTL combination of resistant genotypes exhibited mean survival rates over 90% (see Additional file 1: Table S4), but were rare in the population (< 2%), weakening the confidence in these estimates. These combinations associated Q1.4 with Q1.3 or Q1.5, and Q8.1 with Q28.1. Combination of favorable genotypes for QTL on chromosomes 16 and 1 showed survival rates ranging from 77.2% to 84.0%. The best combination was observed between Q1.2 and Q16.2 while the mean survival rate was only 77.6% for the beneficial combination between Q1.6 and Q16.3 (see Additional file 1: Table S4).

As the combination between Q1.6 and Q16.3 included very promising candidate genes and was also more present in the population than the best combination between Q1.2 and Q16.2 (45% *versus* 24%), we focused in Fig. 3 on the best 3-QTL favorable combinations for the double homozygous resistant genotype (RR-RR) on Q1.6 and Q16.3. We represented all the 3 QTL-combinations exhibiting at least 10% higher mean survival rate for RR-RR-RR than for RR-RR-SS genotypes (Fig. 3). Regarding the mean survival rates of the triple RR combinations, the 3 top combinations (over 85% survival rate) involved Q28.4, Q11.1 and Q28.3 for which the beneficial alleles were the minor ones (Table 3).

**Figure 3.**
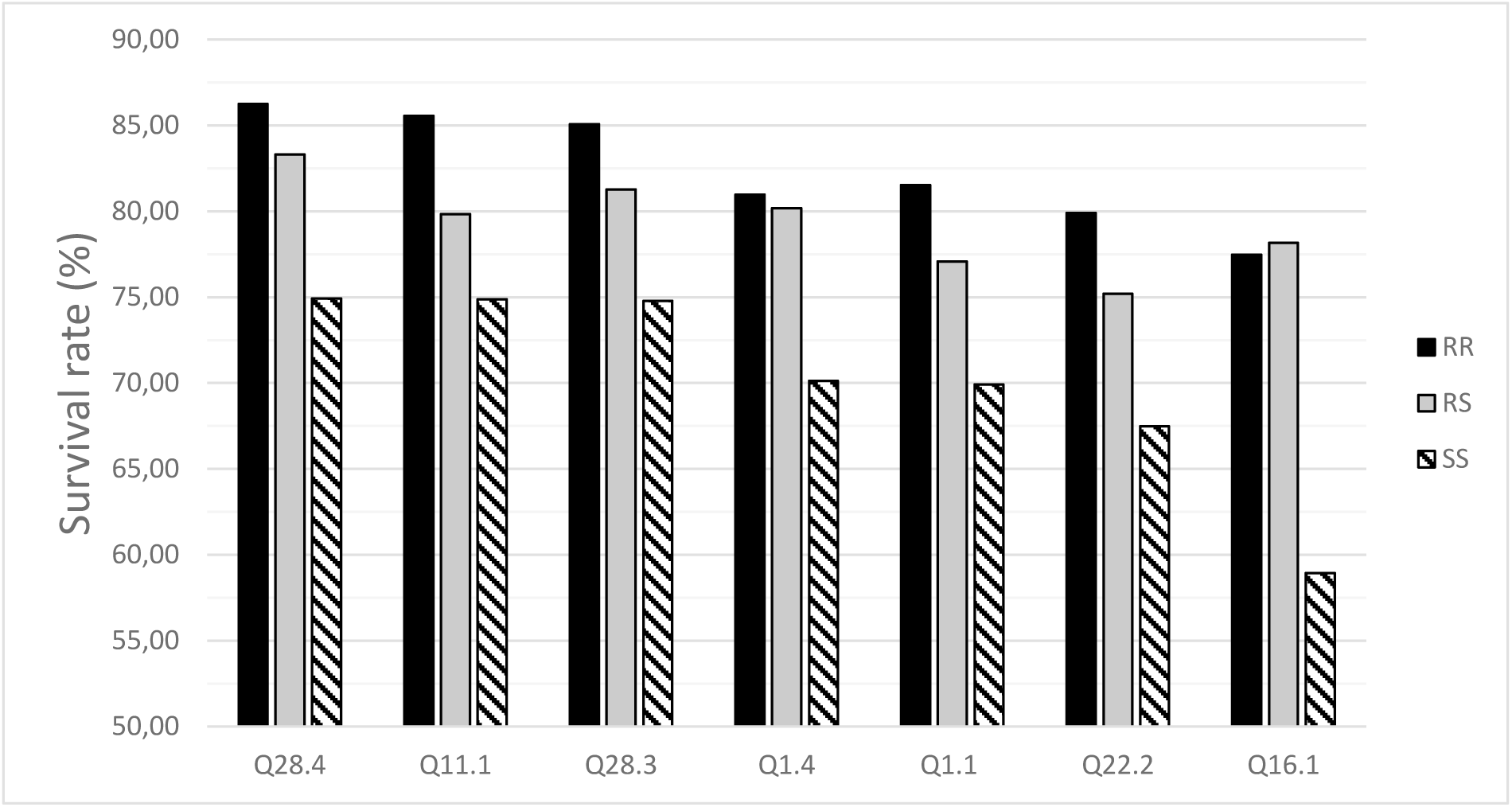
Average survival rate across the two cohorts for the double favorable homozygous on the main QTL Q1.6 (AX-578242608) and Q16.3 (AX-564712030) according to the genotype at the top SNP of a third QTL

## Discussion

The heritability of IPN resistance in Norwegian and Chilean rainbow trout populations was moderate [10,12,43], and our estimation was even a bit lower. These small differences may be explained not only by genetic differences among the French population analyzed here and the two others, but also by important differences in the challenge protocols. First, the challenge was performed in separate tanks for all the 200 families of the Norwegian study [12] while fish from all families were mixed and challenged together in ours and in the chilean study. In addition, the average body weight at challenge was either lower (∼0.2 g; [12]) or higher (∼2.2 g; [10]) than in our study (∼1.5 g). Last but not least, the challenge was by intraperitoneal inoculation in the Chilean case while it was by bath in ours as well as in the Norwegian study. Recently a new estimate (h²=0.21), very close to ours, was derived in a limited full factorial mating design between 5 dams and 5 sires on the Osland Genetics strain, where 610 offspring were bath challenged [44].

Genomic selection is nowadays implemented in the main fish breeding programs because its accuracy is substantially higher than that of traditional pedigree selection based on sib-challenge tests used for assessing disease resistance that cannot be measured directly on selection candidates [45,46]. However, marker-assisted selection (MAS) can have major advantages (reduction of phenotyping and genotyping costs) over traditional or genomic selection for traits whose genetic variation is mainly explained by a few quantitative trait loci (QTL, *i.e* genomic regions harboring genes with a significant effect on the trait). MAS can be applied to quickly eradicate breeders with detrimental genotypes in the selection nucleus, but it can also be used to market some eggs with a high genetic resistance score from multipliers.

Our study confirmed the difference in the number of QTL associated to the resistance to IPNV between the rainbow trout and the Atlantic salmon. While complex polygenic resistance to disease is likely to involve at the level of biological pathways a shared set of genes across populations or close species, however allele frequencies are expected to diverge at loci influencing the host immune response due to natural selection associated to variation in pathogen exposure across populations and geographical areas [47]. It can be hypothesized that the 20 to 30 generations of natural selection of numerous farmed rainbow trout stocks since the first description of an IPN outbreak in 1940 [1] has contributed to generate the actual situation with tens of loci with increased frequencies of beneficial alleles compared to the limited number of Atlantic salmon farmed strains that are currently 10 to 15 generations only from their wild ancestors since the start of breeding programs in the mid-1970s [48]. The major QTL for resistance to IPNV identified in the Atlantic salmon, associated to polymorphic receptor *cdhn1*, has not been found in rainbow trout. Contrary to what has been observed in Atlantic salmon, resistance to IPNV in rainbow trout appears to be polygenic, even though a limited number of SNPs (∼63) explained three-quarters of the additive genetic variance in our French population. Those SNPs were associated to 25 QTL located on 10 out of 32 rainbow trout chromosomes. In particular, 5 QTL exhibited very significant effects with differences in survival rates ranging from over 20% to 30% between homozygous genotypes. Three of these QTL were located on chromosome 1, the 2 others being located on chromosome 16, *i.e.* positioned on the two chromosomes that were previously identified as carrying QTL for resistance to IPNV in rainbow trout [21,25,44,49].

As regards to the QTL previously detected on chromosome 1, Santi et al. [25] indicated in their patent that homozygous fish for the cytosine allele at the most significant SNP AX-89929954 were expected to have a mean survival rate of 45% under conditions similar to those of the challenge test, while homozygous fish for the alternative allele had mean survival rates of 17% and heterozygous fish had a mean survival rates of 36% under similar conditions. The SNP AX-89929954 is located at 18.416812 Mb in the Arlee reference genome (USDA_OmykA_1.1.), the second and third most significant SNPs in [25] being AX-89918280 and AX-89938309 located at 18.738017 Mb and 16.037298 Mb, respectively. All these three positions are located after the end of our last QTL region on chromosome 1 ended at 15.858 Mb, as well as far after the QTL identified in the Osland rainbow trout strain [49] with the top SNP AX-89932951 located at 12.200969 Mb on chromosome 1. Though this last SNP does not belong to the set of quality-filtered SNPs in our analysis, it is worth noting its location in-between our first two QTL on chromosome 1 (Q1.1 and Q1.2, Table 2). However, Ahmad et al. [44] did not confirm any significant association between the SNP AX-89932951 and IPN resistance in a validation cohort of the Osland strain. Differences in the QTL detected among studies may be due to genetic differences across rainbow trout populations, but also to genetic differences in the virus strains. Tapia et al. [50] compared by RNA-seq the transcriptomic response of fry challenged with two Chilean isolates of IPNV, RTTX (genogroup 1), and ALKA (genogroup 5) in rainbow trout. The results revealed that infection with RTTX elicited a most important modulation of the trout transcriptome compared to ALKA infection, generating a greater number of highly differentially expressed genes compared to control fish, especially the first week post-challenge. In the Norwegian studies on rainbow trout [44,49] the virus strain (IPNV-R-L5 strain) used was isolated from Atlantic salmon.

In vitro, in a salmonid cell line, IPNV has been shown to block the type 1 interferon signaling pathway [51] resulting in the limitation of production of the antiviral protein Mx protein particularly effective against IPNV [52]. In Atlantic salmon, even though a clear induction of ISGs could be detected in experimental infections [53,54], asymptomatic carrier state is usually observed in the field [55,56]. The inhibition of both apoptosis and some innate antiviral components is a general hypothesis for IPNV-evading strategies reflected in vitro by the ability of IPNV to persist in cultured cell lines [57,58] and in vivo by the establishment of asymptomatic carrier stage. It is not clear how significant is the carrier stage for the disease control strategies in rainbow trout farming [59,60] but this may be important to evaluate in particular in IPNV resistant selected livestock.

Our GWAS pointed to a number of gene candidates on chromosomes 1, 2, 8, 11, 12, 13, 14, 16, 22 and 28. We identified some QTL on chromosome 8, 11 and 13 where some suggestive QTL were previously detected [44]. However, our top SNPs were at least distant from 8 Mb of the top SNPs identified in their study. In addition, the observed effects of our top SNPs were limited across genotypes (+7 to 13%, Table 3) for these QTL.

In our study, the most interesting QTL were associated to top SNPs with mean survival rate differences over 20% between the beneficial and detrimental genotypes. They were all associated to promising functional candidate genes, and were located on chromosome 1 (*uts2d* (ENSOMYG00000008336, LOC110534233) in Q1.1; *rc3h1* (ENSOMYG00000008605, LOC110488062) in Q1.6; *ga45b* (ENSOMYG00000000138, LOC100301708) in Q1.7), and on chromosome 16 (*irf2bpa* (ENSOMYG00000027318, LOC110491199) in Q16.2; *eif2ak2* (ENSOMYG00000027396, LOC100271898) in Q16.3).

*Urotensin-2* (*uts-2*) has pleiotropic effects on many physiological functions including vasoconstriction, cell division, neuroendocrine activities and inflammation in mammalian models. Pro-inflammatory actions of *uts-2* represent the most direct link with the susceptibility to IPNV [40]. In mammals, UTS-2 stimulates a p38 MAPK dependent pro-inflammatory responses mediated by TNFα, IL1β, IFNγ and IL8 [61]. It also interacts with the IRF3/type I IFN pathway to modulate TNFα and IL1β levels in a p38 MAPK independent manner [62].

The E3 ubiquitin ligase ROQUIN-1 (*rc3h1* gene) has important anti-inflammatory functions and controls T cell responses through decay of targeted mRNAs. ROQUIN protein binds to constitutive decay elements (CDE) or alternative decay elements (ADE) in the 3’ UTR of mRNAs, leading to mRNA deadenylation and degradation. There are CDE in many key genes involved in T cell activation and inflammatory pathways such as *hmgxb3, icos, ier3, nfkbid, nfkbiz, ppp1r10, tnf, tnfrsf4/ox40* [63,64]. Mutations in the mouse gene encoding ROQUIN-1, preventing interaction with its partner REGNASE-1, led to over-activation of T follicular helper cells, abnormal germinal center reaction and auto-antibody production [41].

GADD45 (or GA45) proteins are stress sensors responding to physiological and environmental stimulations, and could affect antiviral responses through modulation of inflammation or via more direct action on antiviral pathways. Indeed, GADD45B has an impact on chemotaxis of inflammatory cells, as reported for gadd45 ^-^/_-_ mice that are unable to recruit granulocytes and macrophages in the peritoneal cavity after injection of bacterial lipopolysaccharides in this location [65]. Through interactions with p38 MAPK and Janus kinase pathways, GADD45 proteins are also important for effectors and inflammatory functions of myeloid cells, such as oxidative burst, cytokine production and phagocytosis [66]. Upon RNA virus infection, GADD45B protein interacts with Ras-GTPase-activating protein (SH3 domain)-binding protein 1 (G3BP1), contributing to stress granule formation and the stress granule-mediated type I IFN response, thereby directly linking these genes to the antiviral response [67]. It should be noticed that all the top SNPs in Q1.6 (associated to *rc3h1*) were in extremely high linkage disequilibrium with the single top SNP defining Q1.7 and located in the *ga45b* gene (Supplementary see Additional file 1: Table S3), suggesting there may be cooperative associations of *rc3h1* and *ga45b* alleles.

Q16.2 is identified by a unique SNP located within the gene *mRNA-interferon regulatory factor 2-binding protein 2-A* (*irf2bp2a*). In mammals, this transcriptional cofactor is involved in different biological systems and plays key roles in both lipid metabolism and control of inflammation [68]. IRF2BP2 specifically down-regulates innate inflammatory response of macrophages, and its expression is strongly repressed during the differentiation of M1 inflammatory macrophages [69]. It is also involved in the regulation of lymphocyte activation [70]. As for genes associated to QTL on chromosome 1, the connexion of *irf2bp2a* to resistance and susceptibility to IPNV would likely be through the modulation of inflammation and/or lymphocyte activation. However, there is no direct evidence about the associated mechanisms involved in immunity against (birna)viruses.

This is in contrast with the gene *eif2ak2* which is associated to Q16.3. *eif2ak2* encodes a dsRNA-dependent serine/threonine-protein kinase (PKR) that phosphorylates the alpha subunit of eukaryotic translation initiation factor 2 (eIF2α). EIF2S1/eIF-2-alpha phosphorylation converts it into a global protein synthesis inhibitor, resulting to a shutdown of cellular and viral protein synthesis, while concomitantly initiating the preferential translation of ISR-specific mRNAs, such as the transcriptional activator ATF4 [71–73]. PKR expression is upregulated by viral infections and type I IFN [74], and it plays a key role in the antiviral innate immune response in fish as well as in mammals (reviewed in [39]). A PKR role in rainbow trout response to IPNV infection has been reported, and *in vitro* loss of function demonstrated that salmonid PKR has conserved molecular functions in apoptosis and translation control [75]. Importantly, treatment with PKR inhibitors led to reduced IPNV titre in CHSE214 cells [76]. Moreover, IPNV infection does not upregulate PKR expression [77], most likely due to a global IPNV strategy of repression of type I IFN response to evade the host antiviral response. In fact, PKR can exert an antiviral activity on a wide range of DNA and RNA viruses including hepatitis C virus (HCV), hepatitis B virus (HBV), measles virus (MV) and herpes simplex virus 1 (HHV-1) [72,78–83]. As an adapter protein and/or via its kinase activity, it triggers multiple signalling pathways including p38 MAPK, NF-kappa-B and insulin signalling pathways), controlling the expression of pro-inflammatory cytokines and IFNs [84–86].

Apart from [44] where AX-89961019 was detected as a suggestive QTL (located at 6.932350 Mb on Arlee reference chromosome 16), only the very first QTL studies on a few Japanese families of rainbow trout detected a QTL on chr16 [20,21], but the few microsatellites used at that time do not allow us to confirm whether or not the same genomic region is detected in our study. Our two main QTL on chr16 (Q16.2 and Q16.3) are located in a ROH island of 2174 kb, previously identified in the LB population [87] between 1.504245 and 3.678601 Mb. This LB population corresponded to an earlier generation of selection of the very same commercial population under our current study. However, this ROH region was not shared with the 3 other populations considered in [87]. ROH island identifies a region of the genome that is frequently homozygous, suggesting a signature of positive selection [88]. This was confirmed as the 3 top SNPs located in Q16.2 and Q16.3 were included in a ROH for 37 to 45% of the fish in the current study (results not shown). The question may arise whether Q16.2 and Q16.3 correspond to a cluster of two genes working together as a supergene [89] to provide an integrated control of a complex response to IPNV.

## Conclusions

Based on functional information collected in the literature, various promising candidate genes were identified, orthologous to genes controlling the type I IFN response or the inflammation response, as well as involved in diverse roles in the cells which might be connected to the virus cycle or to antiviral processes. Remarkably, genes associated to the most significant QTL on chromosomes 1 and 16 are all involved in the regulation of inflammatory pathways such as the p38 MAPK pathway, strongly suggesting a central role of inflammation in the resistance and susceptibility to IPNV in rainbow trout, with PKR being a key factor.

These QTL offer the possibility of marker-assisted selection in rainbow trout for rapid dissemination of genetic improvement for IPN resistance, as it has occurred for the last 15 years in Atlantic salmon, significantly reducing mortality in salmonid aquaculture.

## Declarations

### Ethics approval

Fish experimentation was carried out in strict accordance with European guidelines and recommendations on animal experimentation and welfare (European Directive 2010/63/EU). Experimental procedures were validated by the ANSES animal ethics committee (ANSES/ENVA/UPC No. 16) and authorized by the French Ministry of National Education, Higher Education and Research (APAFIS #2015100516411021 and #2019052112541943).

### Consent for publication

Not applicable

### Availability of data and materials

Please describe where the data can be accessed

### Competing interests

The authors declare that they have no competing interests.

### Funding

This work was supported by the European Maritime and Fisheries Fund and FranceAgrimer (Hypotemp project n° PFEA470019FA1000016, Flavocontrol project n° PFEA470020FA1000004).

### Authors’ contributions

JDA performed the bioinformatic and statistical analyses. AD and SC provided animals for the challenges and contributed to the data acquisition. YF and TM organized the data acquisition and realized the challenges. JDA, YF, PH and FP participated in the design of the study. BC, PB and FP contributed to the interpretation of the analysis. FP provided scientific supervision. FP and PB wrote the first draft of the manuscript. All authors read and approved the final manuscript.

## Acknowledgements

The authors would like to thank (i) the FORTIOR Genetics platform (ANSES, Plouzané, France) for the realization of the challenges, (ii) the INRAE genotyping platform Gentyane (INRAE, Clermont-Ferrand, France) for the production of genotype data. They are also grateful to Pierre Patrice (SYSAAF) for data collection, and Dominique Charles et Nicolas Picchi (Les Aquaculteurs Bretons) for financial and administrative support.

## Additional files

### Additional file 1 Table S1

Format: xlsx

Title: Positions, reference and alternative alleles on the Arlee genome assembly of the 53 top SNPs, as well as the major and minor alleles in the GWAS

### Additional file 1 Table S2

Format: xlsx

Title: Mean survival (%) and genotype frequency (%) for the 2 cohorts G8 and G9 according to the genotypes for the 53 top SNPs

### Additional file 1 Table S3

Format: xlsx

Title: Linkage disequilibrium (r² values) across the 53 top SNPs

### Additional file 1 Table S4

Format: xlsx

Title: Mean survival rates (%) for the benefical homozygotes on a single SNP on the diagonal and for the combination of the beneficial homozygotes for 2 SNPs in the upper triangle with proportion of the combination on the lower triangle

## Notes

### Competing Interest Statement

The authors have declared no competing interest.

## References

[1] M’Gonigle RH. Acute Catarrhal Enteritis of Salmonid Fingerlings. Transactions of the American Fisheries Society 1941;70:297–303. 10.1577/1548-8659(1940)70[297:ACEOSF]2.0.CO;2.

[2] Besse P, Kinkelin PD, Donon O. Sur l’existence en France de la nécrose pancréatique de la truite Arc-en-Ciel (Salmo gairdneri). (Note préliminaire). bavf 1965;118:185–92. 10.4267/2042/67132.

[3] Roberts RJ, Pearson MD. Infectious pancreatic necrosis in Atlantic salmon, *Salmo salar* L. Journal of Fish Diseases 2005;28:383–90. 10.1111/j.1365-2761.2005.00642.x.

[4] Tapia D, Kuznar J, Farlora R, Yáñez JM. Differential Transcriptomic Response of Rainbow Trout to Infection with Two Strains of IPNV. Viruses 2021;14:21. 10.3390/v14010021.

[5] Dopazo CP. The Infectious Pancreatic Necrosis Virus (IPNV) and its Virulence Determinants: What is Known and What Should be Known. Pathogens 2020;9:94. 10.3390/pathogens9020094.

[6] Duan K, Tang X, Zhao J, Ren G, Shao Y, Lu T, et al. An inactivated vaccine against infectious pancreatic necrosis virus in rainbow trout (Oncorhynchus mykiss). Fish & Shellfish Immunology 2022;127:48–55. 10.1016/j.fsi.2022.06.008.

[7] Li L, Liu W, Zhang Z, Zhao J, Lu T, Shao Y, et al. IPNV inactive vaccine supplemented with GEL 02 PR adjuvant: Protective efficacy, cross-protection, and stability. Fish & Shellfish Immunology 2025;158:110167. 10.1016/j.fsi.2025.110167.

[8] Li S, Li X, Yuan R, Chen X, Chen S, Qiu Y, et al. Development of a recombinant adenovirus-vectored vaccine against both infectious hematopoietic necrosis virus and infectious pancreatic necrosis virus in rainbow trout (Oncorhynchus mykiss). Fish & Shellfish Immunology 2023;132:108457. 10.1016/j.fsi.2022.108457.

[9] Yáñez JM, Houston RD, Newman S. Genetics and genomics of disease resistance in salmonid species. Front Genet 2014;5. 10.3389/fgene.2014.00415.

[10] Flores-Mara R, Rodríguez FH, Bangera R, Lhorente JP, Neira R, Newman S, et al. Resistance against infectious pancreatic necrosis exhibits significant genetic variation and is not genetically correlated with harvest weight in rainbow trout (Oncorhynchus mykiss). Aquaculture 2017;479:155–60. 10.1016/j.aquaculture.2017.05.042.

[11] Rodríguez FH, Flores-Mara R, Yoshida GM, Barría A, Jedlicki AM, Lhorente JP, et al. Genome-Wide Association Analysis for Resistance to Infectious Pancreatic Necrosis Virus Identifies Candidate Genes Involved in Viral Replication and Immune Response in Rainbow Trout (*Oncorhynchus mykiss*). G3 Genes|Genomes|Genetics 2019;9:2897–904. 10.1534/g3.119.400463.

[12] Wetten M, Kjøglum S, Fjalestad KT, Skjaervik O, Storset A. Genetic variation in resistance to infectious pancreatic necrosis in rainbow trout (Oncorhynchus mykiss) after a challenge test: Breeding for IPN-resistance in rainbow trout. Aquaculture Research 2011;42:1745–51. 10.1111/j.1365-2109.2010.02771.x.

[13] Houston RD, Haley CS, Hamilton A, Guy DR, Tinch AE, Taggart JB, et al. Major Quantitative Trait Loci Affect Resistance to Infectious Pancreatic Necrosis in Atlantic Salmon (*Salmo salar*). Genetics 2008;178:1109–15. 10.1534/genetics.107.082974.

[14] Moen T, Baranski M, Sonesson AK, Kjøglum S. Confirmation and fine-mapping of a major QTL for resistance to infectious pancreatic necrosis in Atlantic salmon (Salmo salar): population-level associations between markers and trait. BMC Genomics 2009;10:368. 10.1186/1471-2164-10-368.

[15] Houston RD, Haley CS, Hamilton A, Guy DR, Mota-Velasco JC, Gheyas AA, et al. The susceptibility of Atlantic salmon fry to freshwater infectious pancreatic necrosis is largely explained by a major QTL. Heredity 2010;105:318–27. 10.1038/hdy.2009.171.

[16] Moen T, Torgersen J, Santi N, Davidson WS, Baranski M, Ødegård J, et al. Epithelial Cadherin Determines Resistance to Infectious Pancreatic Necrosis Virus in Atlantic Salmon. Genetics 2015;200:1313–26. 10.1534/genetics.115.175406.

[17] Bishop SC, Woolliams JA. Genomics and disease resistance studies in livestock. Livestock Science 2014;166:190–8. 10.1016/j.livsci.2014.04.034.

[18] Moen T, Ødegård J. Genomics in Selective Breeding of Atlantic Salmon, Vancouver, BC, Canada.: 2014.

[19] Hillestad B, Johannessen S, Melingen GO, Moghadam HK. Identification of a New Infectious Pancreatic Necrosis Virus (IPNV) Variant in Atlantic Salmon (Salmo salar L.) that can Cause High Mortality Even in Genetically Resistant Fish. Front Genet 2021;12:635185. 10.3389/fgene.2021.635185.

[20] Ozaki A, Khoo S-K, Yoshiura Y, Ototake M, Sakamoto T, Dijkstra JM, et al. Identification of Additional Quantitative Trait Loci (QTL) Responsible for Susceptibility to Infectious Pancreatic Necrosis Virus in Rainbow Trout. Fish Pathol 2007;42:131–40. 10.3147/jsfp.42.131.

[21] Ozaki A, Sakamoto T, Khoo S, Nakamura K, Coimbra MR, Akutsu T, et al. Quantitative trait loci (QTLs) associated with resistance/susceptibility to infectious pancreatic necrosis virus (IPNV) in rainbow trout (Oncorhynchus mykiss). Mol Gen Genomics 2001;265:23–31. 10.1007/s004380000392.

[22] Phillips RB, Nichols KM, DeKoning JJ, Morasch MR, Keatley KA, Rexroad C, et al. Assignment of Rainbow Trout Linkage Groups to Specific Chromosomes. Genetics 2006;174:1661–70. 10.1534/genetics.105.055269.

[23] Danzmann RG, Cairney M, Davidson WS, Ferguson MM, Gharbi K, Guyomard R, et al. A comparative analysis of the rainbow trout genome with 2 other species of fish (Arctic charr and Atlantic salmon) within the tetraploid derivative Salmonidae family (subfamily: Salmoninae). Genome 2005;48:1037–51. 10.1139/g05-067.

[24] Pearse DE, Barson NJ, Nome T, Gao G, Campbell MA, Abadía-Cardoso A, et al. Sex-dependent Dominance Maintains Migration Supergene in Rainbow Trout. Nat. Ecol. Evol. 2019;3:1731–42. 10.1038/s41559-019-1076-y.

[25] Santi N, Moen T, Ødegård J. METHOD FOR PREDICTING RESISTANCE. US20190241980A1, 2019.

[26] Gao G, Magadan S, Waldbieser GC, Youngblood RC, Wheeler PA, Scheffler BE, et al. A long reads-based *de-novo* assembly of the genome of the Arlee homozygous line reveals chromosomal rearrangements in rainbow trout. G3 Genes|Genomes|Genetics 2021;11:jkab052. 10.1093/g3journal/jkab052.

[27] Bernard M, Dehaullon A, Gao G, Paul K, Lagarde H, Charles M, et al. Development of a High-Density 665 K SNP Array for Rainbow Trout Genome-Wide Genotyping. Front Genet 2022;13:941340. 10.3389/fgene.2022.941340.

[28] Chang CC, Chow CC, Tellier LC, Vattikuti S, Purcell SM, Lee JJ. Second-generation PLINK: rising to the challenge of larger and richer datasets. Gigascience 2015;4:s13742-015-0047–8. 10.1186/s13742-015-0047-8.

[29] Purcell S, Neale B, Todd-Brown K, Thomas L, Ferreira MAR, Bender D, et al. PLINK: A Tool Set for Whole-Genome Association and Population-Based Linkage Analyses. The American Journal of Human Genetics 2007;81:559–75. 10.1086/519795.

[30] Griot R, Allal F, Brard-Fudulea S, Morvezen R, Haffray P, Phocas F, et al. APIS: An auto-adaptive parentage inference software that tolerates missing parents. Molecular Ecology Resources 2020;20:579–90. 10.1111/1755-0998.13103.

[31] Roche J, Griot R, Besson M, Allal F, Vandeputte M, D’Ambrosio J, et al. APIS: Auto-Adaptive Parentage Inference Software Tolerant to Missing Parents 2024.

[32] Sargolzaei M, Chesnais JP, Schenkel FS. A new approach for efficient genotype imputation using information from relatives. BMC Genomics 2014;15:478. 10.1186/1471-2164-15-478.

[33] Misztal I, Tsuruta S, Lourenco D, Masuda Y, Aguilar I, Legarra A, et al. Manual for BLUPF90 family of programs. url : http://nce.ads.uga.edu/wiki/lib/exe/fetch.php?media=blupf90_all7.pdf. 2014.

[34] VanRaden PM. Efficient Methods to Compute Genomic Predictions. Journal of Dairy Science 2008;91:4414–23. 10.3168/jds.2007-0980.

[35] Zhou X, Carbonetto P, Stephens M. Polygenic Modeling with Bayesian Sparse Linear Mixed Models. PLoS Genet 2013;9:e1003264. 10.1371/journal.pgen.1003264.

[36] Stephens M, Balding DJ. Bayesian statistical methods for genetic association studies. Nat Rev Genet 2009;10:681–90. 10.1038/nrg2615.

[37] Kass RE, and Raftery AE. Bayes Factors. Journal of the American Statistical Association 1995;90:773–95. 10.1080/01621459.1995.10476572.

[38] Michenet A, Saintilan R, Venot E, Phocas F. Insights into the genetic variation of maternal behavior and suckling performance of continental beef cows. Genet Sel Evol 2016;48:45. 10.1186/s12711-016-0223-z.

[39] Chaumont L, Collet B, Boudinot P. Double-stranded RNA-dependent protein kinase (PKR) in antiviral defence in fish and mammals. Developmental & Comparative Immunology 2023;145:104732. 10.1016/j.dci.2023.104732.

[40] Sun S, Liu L. Urotensin II: an inflammatory cytokine. Journal of Endocrinology 2019;240:R107–17. 10.1530/JOE-18-0505.

[41] Behrens G, Edelmann SL, Raj T, Kronbeck N, Monecke T, Davydova E, et al. Disrupting Roquin-1 interaction with Regnase-1 induces autoimmunity and enhances antitumor responses. Nat Immunol 2021;22:1563–76. 10.1038/s41590-021-01064-3.

[42] Essig K, Kronbeck N, Guimaraes JC, Lohs C, Schlundt A, Hoffmann A, et al. Roquin targets mRNAs in a 3′-UTR-specific manner by different modes of regulation. Nat Commun 2018;9:3810. 10.1038/s41467-018-06184-3.

[43] Yoshida GM, Carvalheiro R, Rodríguez FH, Lhorente JP, Yáñez JM. Single-step genomic evaluation improves accuracy of breeding value predictions for resistance to infectious pancreatic necrosis virus in rainbow trout. Genomics 2019;111:127–32. 10.1016/j.ygeno.2018.01.008.

[44] Ahmad A, Aslam ML, Evensen Ø, Gamil AAA, Berge A, Solberg T, et al. The genetics of resistance to infectious pancreatic necrosis virus in rainbow trout unveiled through survival and virus load data. Front Genet 2024;15:1484287. 10.3389/fgene.2024.1484287.

[45] Sonesson AK, Meuwissen TH. Testing strategies for genomic selection in aquaculture breeding programs. Genet Sel Evol 2009;41:37. 10.1186/1297-9686-41-37.

[46] Villanueva B, Fernández J, García-Cortés LA, Varona L, Daetwyler HD, Toro MA. Accuracy of genome-wide evaluation for disease resistance in aquaculture breeding programs1. Journal of Animal Science 2011;89:3433–42. 10.2527/jas.2010-3814.

[47] Randolph HE, Aracena KA, Lin Y, Mu Z, Barreiro LB. Shaping immunity: The influence of natural selection on population immune diversity. Immunological Reviews 2024;323:227–40. 10.1111/imr.13329.

[48] Gjedrem T. Genetic improvement for the development of efficient global aquaculture: A personal opinion review. Aquaculture 2012;344–349:12–22. 10.1016/j.aquaculture.2012.03.003.

[49] Aslam M, Valdemarsson S, Berg A, Gjerde B. Book of abstracts of the 70th annual meeting of the European federation of animal science 2019.

[50] Tapia D, Eissler Y, Reyes-Lopez FE, Kuznar J, Yáñez JM. Infectious pancreatic necrosis virus in salmonids: Molecular epidemiology and host response to infection. Reviews in Aquaculture 2022;14:751–69. 10.1111/raq.12623.

[51] Collet B. Innate immune responses of salmonid fish to viral infections. Developmental & Comparative Immunology 2014;43:160–73. 10.1016/j.dci.2013.08.017.

[52] Lester K, Hall M, Urquhart K, Gahlawat S, Collet B. Development of an in vitro system to measure the sensitivity to the antiviral Mx protein of fish viruses. Journal of Virological Methods 2012;182:1–8. 10.1016/j.jviromet.2012.01.014.

[53] Ellis AE, Cavaco A, Petrie A, Lockhart K, Snow M, Collet B. Histology, immunocytochemistry and qRT-PCR analysis of Atlantic salmon, Salmo salar L., post-smolts following infection with infectious pancreatic necrosis virus (IPNV). Journal of Fish Diseases 2010;33:803–18. 10.1111/j.1365-2761.2010.01174.x.

[54] Mcbeath A, Snow M, Secombes C, Ellis A, Collet B. Expression kinetics of interferon and interferon-induced genes in Atlantic salmon (Salmo salar) following infection with infectious pancreatic necrosis virus and infectious salmon anaemia virus. Fish & Shellfish Immunology 2007;22:230–41. 10.1016/j.fsi.2006.05.004.

[55] Lockhart K, Gahlawat SK, Soto-Mosquera D, Bowden TJ, Ellis AE. IPNV carrier Atlantic salmon growers do not express Mx mRNA and poly I:C-induced Mx response does not cure the carrier state. Fish & Shellfish Immunology 2004;17:347–52. 10.1016/j.fsi.2004.04.011.

[56] Ørpetveit I, Mikalsen AB, Sindre H, Evensen Ø, Dannevig BH, Midtlyng PJ. Detection of *Infectious Pancreatic Necrosis Virus* in Subclinically Infected Atlantic Salmon by Virus Isolation in Cell Culture or Real-Time Reverse Transcription Polymerase Chain Reaction: Influence of Sample Preservation and Storage. J VET Diagn Invest 2010;22:886–95. 10.1177/104063871002200606.

[57] Jurado MT, García-Valtanen P, Estepa A, Perez L. Antiviral activity produced by an IPNV-carrier EPC cell culture confers resistance to VHSV infection. Veterinary Microbiology 2013;166:412–8. 10.1016/j.vetmic.2013.06.022.

[58] Rodríguez Saint-Jean S, De Las Heras AI, Pérez Prieto SI. The persistence of infectious pancreatic necrosis virus and its influence on the early immune response. Veterinary Immunology and Immunopathology 2010;136:81–91. 10.1016/j.vetimm.2010.02.015.

[59] Ahne W, Thomsen I. Infectious Pancreatic Necrosis: Detection of Virus and Antibodies in Rainbow Trout IPNV-Carrier (Salmo gairdneri). Journal of Veterinary Medicine, Series B 1986;33:552–4. 10.1111/j.1439-0450.1986.tb00067.x.

[60] Rodriguez S, Alonso M, Perez-Prieto SI. Detection of Infections Pancreatic Necrosis Virus(IPNV) from Leukocytes of Carrier Rainbow Trout Oncorhynchus mykiss. Fish Pathol 2001;36:139–46. 10.3147/jsfp.36.139.

[61] Liu LM, Liang DY, Ye CG, Tu WJ, Zhu T. The UII/UT System Mediates Upregulation of Proinflammatory Cytokines through p38 MAPK and NF-κB Pathways in LPS-Stimulated Kupffer Cells. PLoS ONE 2015;10:e0121383. 10.1371/journal.pone.0121383.

[62] Liu L, Tu W, Zhu T, Wang X, Tan Z, Zhong H, et al. IRF3 is an important molecule in the UII/UT system and mediates immune inflammatory injury in acute liver failure. Oncotarget 2016;7:49027–41. 10.18632/oncotarget.10717.

[63] Tan D, Zhou M, Kiledjian M, Tong L. The ROQ domain of Roquin recognizes mRNA constitutive-decay element and double-stranded RNA. Nature Structural & Molecular Biology 2014;21:679–85. 10.1038/nsmb.2857.

[64] Tavernier SJ, Athanasopoulos V, Verloo P, Behrens G, Staal J, Bogaert DJ, et al. Author Correction: A human immune dysregulation syndrome characterized by severe hyperinflammation with a homozygous nonsense Roquin-1 mutation. Nat Commun 2019;10:5337. 10.1038/s41467-019-13379-9.

[65] Salerno DM, Tront JS, Hoffman B, Liebermann DA. Gadd45a and Gadd45b modulate innate immune functions of granulocytes and macrophages by differential regulation of p38 and JNK signaling. Journal Cellular Physiology 2012;227:3613–20. 10.1002/jcp.24067.

[66] Hoffman B, Liebermann DA. Gadd45 modulation of intrinsic and extrinsic stress responses in myeloid cells. Journal Cellular Physiology 2009;218:26–31. 10.1002/jcp.21582.

[67] Chathuranga WAG, Nikapitiya C, Kim J-H, Chathuranga K, Weerawardhana A, Dodantenna N, et al. Gadd45β is critical for regulation of type I interferon signaling by facilitating G3BP-mediated stress granule formation. Cell Reports 2023;42:113358. 10.1016/j.celrep.2023.113358.

[68] Chen H-H, Keyhanian K, Zhou X, Vilmundarson RO, Almontashiri NAM, Cruz SA, et al. IRF2BP2 Reduces Macrophage Inflammation and Susceptibility to Atherosclerosis. Circulation Research 2015;117:671–83. 10.1161/CIRCRESAHA.114.305777.

[69] Zhang H, Reilly MP. IRF2BP2: A New Player at the Crossroads of Inflammation and Lipid Metabolism. Circulation Research 2015;117:656–8. 10.1161/CIRCRESAHA.115.307245.

[70] Ramalho-Oliveira R, Oliveira-Vieira B, Viola JPB. IRF2BP2: A new player in the regulation of cell homeostasis. Journal of Leukocyte Biology 2019;106:717–23. 10.1002/JLB.MR1218-507R.

[71] Harashima A, Guettouche T, Barber GN. Phosphorylation of the NFAR proteins by the dsRNA-dependent protein kinase PKR constitutes a novel mechanism of translational regulation and cellular defense. Genes Dev 2010;24:2640–53. 10.1101/gad.1965010.

[72] Kang J-I, Kwon S-N, Park S-H, Kim YK, Choi S-Y, Kim JP, et al. PKR protein kinase is activated by hepatitis C virus and inhibits viral replication through translational control. Virus Research 2009;142:51–6. 10.1016/j.virusres.2009.01.007.

[73] Okumura F, Okumura AJ, Uematsu K, Hatakeyama S, Zhang D-E, Kamura T. Activation of Double-stranded RNA-activated Protein Kinase (PKR) by Interferon-stimulated Gene 15 (ISG15) Modification Down-regulates Protein Translation. Journal of Biological Chemistry 2013;288:2839–47. 10.1074/jbc.M112.401851.

[74] Meurs E, Chong K, Galabru J, Thomas NSB, Kerr IM, Williams BRG, et al. Molecular cloning and characterization of the human double-stranded RNA-activated protein kinase induced by interferon. Cell 1990;62:379–90. 10.1016/0092-8674(90)90374-N.

[75] Chaumont L, Peruzzi M, Huetz F, Raffy C, Le Hir J, Minke J, et al. Salmonid Double-stranded RNA–Dependent Protein Kinase Activates Apoptosis and Inhibits Protein Synthesis. The Journal of Immunology 2024;213:700–17. 10.4049/jimmunol.2400076.

[76] Gamil A, Xu C, Mutoloki S, Evensen Ø. PKR Activation Favors Infectious Pancreatic Necrosis Virus Replication in Infected Cells. Viruses 2016;8:173. 10.3390/v8060173.

[77] Gamil A, Mutoloki S, Evensen Ø. A Piscine Birnavirus Induces Inhibition of Protein Synthesis in CHSE-214 Cells Primarily through the Induction of eIF2α Phosphorylation. Viruses 2015;7:1987–2005. 10.3390/v7041987.

[78] Cassady KA, Gross M. The Herpes Simplex Virus Type 1 U_S_ 11 Protein Interacts with Protein Kinase R in Infected Cells and Requires a 30-Amino-Acid Sequence Adjacent to a Kinase Substrate Domain. J Virol 2002;76:2029–35. 10.1128/jvi.76.5.2029-2035.2002.

[79] Chang J-H, Kato N, Muroyama R, Taniguchi H, Guleng B, Dharel N, et al. Double-stranded RNA-activated protein kinase inhibits hepatitis C virus replication but may be not essential in interferon treatment. Liver International 2010;30:311–8. 10.1111/j.1478-3231.2009.02144.x.

[80] Lin SS, Lee DCW, Law AHY, Fang JW, Chua DTT, Lau ASY. A role for protein kinase PKR in the mediation of Epstein–Barr virus latent membrane protein-1-induced IL-6 and IL-10 expression. Cytokine 2010;50:210–9. 10.1016/j.cyto.2010.01.008.

[81] Okonski KM, Samuel CE. Stress Granule Formation Induced by Measles Virus Is Protein Kinase PKR Dependent and Impaired by RNA Adenosine Deaminase ADAR1. J Virol 2013;87:756–66. 10.1128/JVI.02270-12.

[82] Park I-H, Baek K-W, Cho E-Y, Ahn B-Y. PKR-Dependent Mechanisms of Interferon-α for Inhibiting Hepatitis B Virus Replication. Molecules and Cells 2011;32:167–72. 10.1007/s10059-011-1059-6.

[83] Zhang L, Alter HJ, Wang H, Jia S, Wang E, Marincola FM, et al. The modulation of hepatitis C virus 1a replication by PKR is dependent on NF-kB mediated interferon beta response in Huh7.5.1 cells. Virology 2013;438:28–36. 10.1016/j.virol.2013.01.015.

[84] Li Y, Xie J, Wu S, Xia J, Zhang P, Liu C, et al. Protein Kinase Regulated by dsRNA Downregulates the Interferon Production in Dengue Virus- and dsRNA-Stimulated Human Lung Epithelial Cells. PLoS ONE 2013;8:e55108. 10.1371/journal.pone.0055108.

[85] McAllister CS, Taghavi N, Samuel CE. Protein Kinase PKR Amplification of Interferon β Induction Occurs through Initiation Factor eIF-2α-mediated Translational Control. Journal of Biological Chemistry 2012;287:36384–92. 10.1074/jbc.M112.390039.

[86] Shen S, Niso-Santano M, Adjemian S, Takehara T, Malik SA, Minoux H, et al. Cytoplasmic STAT3 Represses Autophagy by Inhibiting PKR Activity. Molecular Cell 2012;48:667–80. 10.1016/j.molcel.2012.09.013.

[87] Paul K, Restoux G, Phocas F. Genome-wide detection of positive and balancing signatures of selection shared by four domesticated rainbow trout populations (Oncorhynchus mykiss). Genet Sel Evol 2024;56:13. 10.1186/s12711-024-00884-9.

[88] Saravanan KA, Panigrahi M, Kumar H, Parida S, Bhushan B, Gaur GK, et al. Genomic scans for selection signatures revealed candidate genes for adaptation and production traits in a variety of cattle breeds. Genomics 2021;113:955–63. 10.1016/j.ygeno.2021.02.009.

[89] Schwander T, Libbrecht R, Keller L. Supergenes and Complex Phenotypes. Current Biology 2014;24:R288–94. 10.1016/j.cub.2014.01.056.

